# Longitudinal expression profiling of CD4+ and CD8+ cells in patients with active to quiescent Giant Cell Arteritis

**DOI:** 10.1101/243493

**Authors:** Elisabeth De Smit, Samuel W Lukowski, Lisa Anderson, Anne Senabouth, Kaisar Dauyey, Sharon Song, Bruce Wyse, Lawrie Wheeler, Christine Y Chen, Khoa Cao, Amy Wong Ten Yuen, Neil Shuey, Linda Clarke, Isabel Lopez Sanchez, Sandy SC Hung, Alice Pébay, David A Mackey, Matthew A Brown, Alex W Hewitt, Joseph E Powell

**Affiliations:** Centre for Eye Research Australia, The University of Melbourne, Royal Victorian Eye & Ear Hospital, East Melbourne, AUSTRALIA; Institute for Molecular Bioscience, University of Queensland, Brisbane, QLD, AUSTRALIA; Institute of Health and Biomedical Innovation, Queensland University of Technology, Translational Research Institute, Princess Alexandra Hospital, Brisbane, QLD, AUSTRALIA; Ophthalmology Department at Monash Health, Department of Surgery, School of Clinical Sciences at Monash Health, VIC, AUSTRALIA; Department of Neuro-Ophthalmology, Royal Victorian Eye and Ear Hospital, VIC, AUSTRALIA; Centre for Ophthalmology and Visual Science, University of Western Australia, Lions Eye Institute, Perth, AUSTRALIA; School of Medicine, Menzies Research Institute Tasmania, University of Tasmania, Hobart, AUSTRALIA

## Abstract

**Background:** Giant cell arteritis (GCA) is the most common form of vasculitis affecting elderly people. It is one of the few true ophthalmic emergencies. GCA is a heterogenous disease, symptoms and signs are variable thereby making it challenging to diagnose and often delaying diagnosis. A temporal artery biopsy is the gold standard to test for GCA, and there are currently no specific biochemical markers to categorize or aid diagnosis of the disease. We aimed to identify a less invasive method to confirm the diagnosis of GCA, as well as to ascertain clinically relevant predictive biomarkers by studying the transcriptome of purified peripheral CD4+ and CD8+ T lymphocytes in patients with GCA.

**Methods and Findings:** We recruited 16 patients with histological evidence of GCA at the Royal Victorian Eye and Ear Hospital (RVEEH), Melbourne, Australia, and aimed to collect blood samples at six time points: acute phase, 2–3 weeks, 6–8 weeks, 3 months, 6 months and 12 months after clinical diagnosis. CD4+ and CD8+ T-cells were positively selected at each time point through magnetic-assisted cell sorting (MACS). RNA was extracted from all 195 collected samples for subsequent RNA sequencing. The expression profiles of patients were compared to those of 16 age-matched controls. Over the 12-month study period, polynomial modelling analyses identified 179 and 4 statistically significant transcripts with altered expression profiles (FDR < 0.05) between cases and controls in CD4+ and CD8+ populations, respectively. In CD8+ cells, we identified two transcripts that remained differentially expressed after 12 months, namely *SGTB*, associated with neuronal apoptosis, and *FCGR3A*, which has been found in association with Takayasu arteritis (TA), another large vessel vasculitis. We detected genes that correlate with both symptoms and biochemical markers used in the acute setting for predicting long-term prognosis. 15 genes were shared across 3 phenotypes in CD4 and 16 across CD8 cells. In CD8, *IL32* was common to 5 phenotypes: a history of Polymyalgia Rheumatica, both visual disturbance and raised neutrophils at the time of presentation, bilateral blindness and death within 12 months. Altered *IL32* gene expression could provide risk evaluation of GCA diagnosis at the time of presentation and give an indication of prognosis, which may influence management.

**Conclusions:** This is the first longitudinal gene expression study undertaken to identify robust transcriptomic biomarkers of GCA. Our results show cell type-specific transcript expression profiles, novel gene-phenotype associations, and uncover important biological pathways for this disease. These data significantly enhance the current knowledge of relevant biomarkers, their association with clinical prognostic markers, as well as potential candidates for detecting disease activity in whole blood samples. In the acute phase, the gene-phenotype relationships we have identified could provide insight to potential disease severity and as such guide us in initiating appropriate patient management.

## INTRODUCTION

Giant Cell Arteritis (GCA) is the most common form of vasculitis in people over 50 years of age, and has a predilection for medium- and large-sized vessels of the head and neck. GCA represents one of the few true ophthalmic emergencies, and given the severe sequelae of untreated disease, a timely diagnosis is crucial [1]. GCA is a devastating disease associated with significant morbidity and mortality. If untreated, GCA can cause catastrophic complications including blindness and stroke, as well as aortic dissection and rupture.

The patho-aetiology of GCA is poorly understood. It is likely that both a genetic predisposition and possible environmental factors, the latter unconfirmed, contribute to the onset of disease [2]. GCA is a heterogenous disease and a definitive diagnosis can be difficult to establish in the acute setting. The current gold standard for diagnosis is a temporal artery biopsy, which is an invasive surgical procedure [3,4]. There are currently no specific biomarkers to diagnose GCA, or stratify patient management.

In the acute setting, treatment with high-dose corticosteroids should be started empirically when a patient’s symptoms and/or inflammatory markers suggest a diagnosis of GCA is likely [1]. Treatment should not be delayed whilst waiting for biopsy results to become available. Once diagnosed, clinicians monitor disease activity based on patients’ symptoms and inflammatory markers, primarily the erythrocyte sedimentation rate (ESR) and C-reactive protein (CRP). However, these biochemical markers are nonspecific and may be elevated in other inflammatory or infective diagnoses. There is a pressing need for more sensitive and specific biomarkers. This would aid in making a diagnosis, as well managing this condition more appropriately and mitigate the need for an invasive surgical procedure. Motivated by this need, we aimed to discover a biomarker so that when patients present to the emergency department with features of GCA, a blood test could be performed, allowing prompt diagnosis and initiation of appropriate treatment.

GCA is presumed to be an autoimmune disease with a highly complex immunopathogenesis. It has a strong association with HLA class II suggesting an adaptive immune response with antigen presentation to CD4+ T cells [5]. CD8+ T cells have also been described in GCA both at tissue level and peripherally [6,7]. Transcriptional profiling in blood consists of measuring RNA abundance in circulating nucleated cells. Changes in transcript abundance can result from exposure to host- or pathogen-derived immunogenic factors. Given that T Lymphocytes are key mediators of the adaptive cellular immune response and in GCA [8], we studied the transcriptome of peripheral CD4+ and CD8+ T cells of patients with GCA. We monitored patients’ expression profiling along the course of their disease to detect changes in transcripts as disease state altered and became quiescent.

## METHODS

### Patient recruitment

Between July 2014 and June 2016, 16 patients presenting to the emergency department (ED) at the Royal Victorian Eye & Ear Hospital (RVEEH) in Melbourne (Australia), with symptoms and signs consistent with the diagnosis of GCA were enrolled in our study (Figure 1). Ethics was approved for this study through the RVEEH (Ethics 11/998H), and all patients provided informed written consent to participate in serial sample collections. We acquired blood samples from patients in the acute phase of their disease T1 (Day 0–7) but ideally prior to steroid initiation. Analysis took into account those patients who were steroid-naive at T1 and those who had already started steroid treatment, albeit in some cases less than 24 hours earlier. In addition to T1, we aimed to acquire five subsequent serial samples from each patient - T2 (2–3 weeks), T3 (6–8 weeks), T4 (~3 months), T5 (~6 months) and T6 (~12 months) after presentation - to detect changes in their transcripts as the disease state altered and became quiescent (Supplementary Table 1). For each patient with GCA, we recruited an age- and gender-matched healthy control from whom two serial blood samples were collected 2–3 weeks apart. Our study design is outlined in Figure 1.

**Figure 1.**
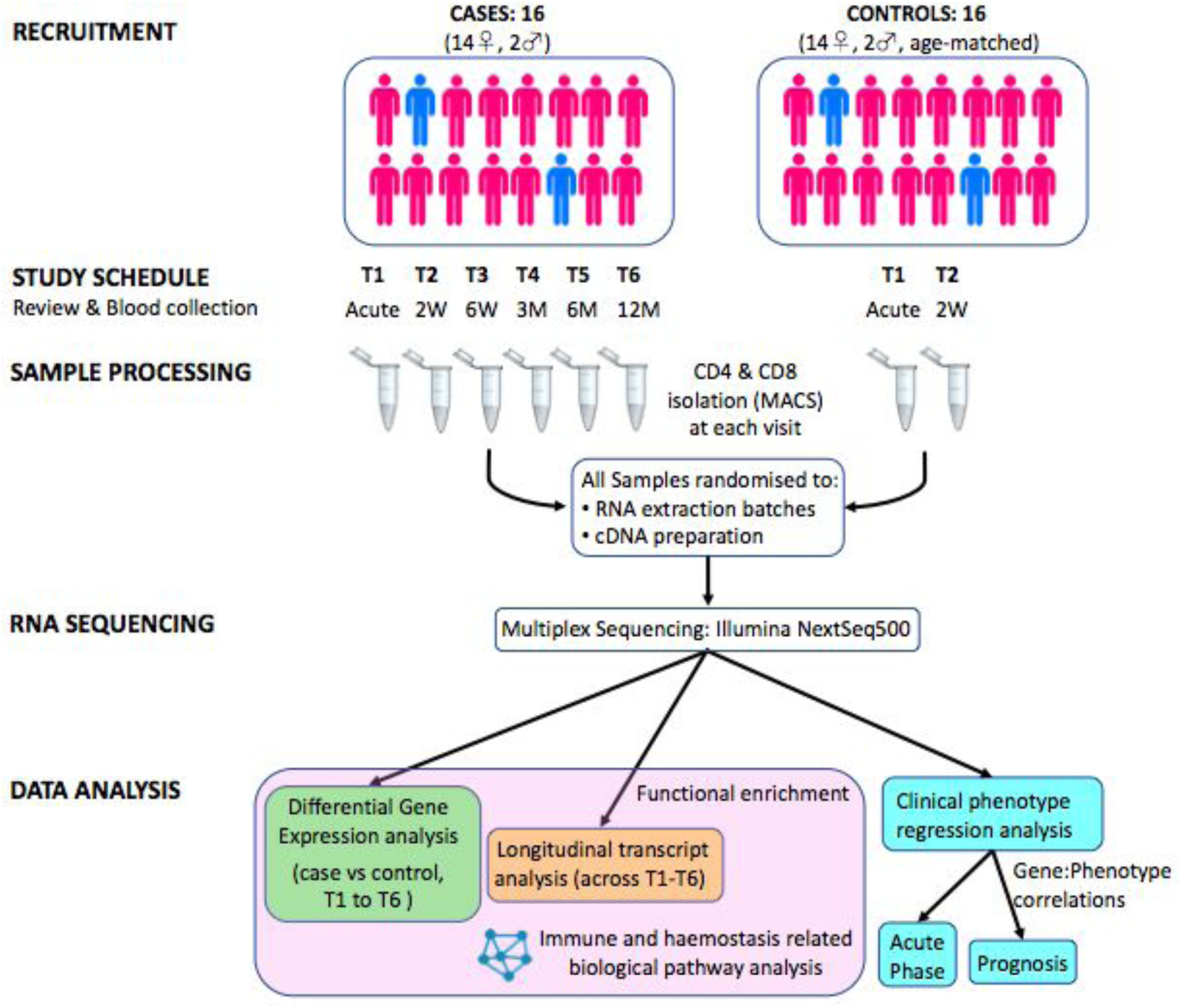
Overview of the study design. A total of 16 patients with GCA had serial blood tests to investigate the gene expression profiles of T lymphocytes over the course of their disease. CD4+ and CD8+ cells were positively selected through magnetic assisted cell sorting (MACS). RNA was extracted for subsequent RNA sequencing. The expression profiles of patients were compared to that of 16 age-matched controls. In addition to differential gene expression analysis and longitudinal transcript analysis, clinical phenotype regression analysis was performed to investigate genes predictive of acute disease and prognosis.

### T-cell isolation

At each visit, 36 ml of peripheral blood were collected in 4 × 9 ml ethylenediaminetetraacetic acid (EDTA) tubes, 18 ml of which were used to isolate each of the two T-cell populations. Once blood was collected from a patient, it was processed within 30 minutes. Rapid processing was conducted to avoid changes in cellular expression profiles [9]. First, the peripheral blood mononuclear cells (PBMCs) were isolated using Ficoll-Paque density centrifugation. This was followed by positive selection with magnetic antibody-coupled microbeads (MACS) (CD4 Human Microbeads (130–045–101) and CD8 Human Microbeads (130–045–201) from Miltenyi Biotec), to isolate the CD4+ and CD8+ T-cell populations from PBMCs. CD4+ cells were labelled with fluorescein isothiocyanate (CD4-Viobright FITC (130–104–515) Miltenyi Biotec) and CD8+ with allophycocyanin (CD8-APC (130–091–076) Miltenyi Biotec) antibody for purity analysis. The CD4+ and CD8+ positive fractions were eluted from the magnetically charged MS column in 1000ul of MACS BSA Stock Solution 1:20 with autoMACS Rinsing Solution (Miltenyi Biotec). A 20 μl aliquot of both CD4+ and CD8+ final cell populations was fixed in 2% paraformaldehyde (PFA) and used for analysis of the population purity on a CyAn ADP fluorescence-activated cell sorting (FACS) analyzer (Supplementary Figure 1). The remainder of the positive fractions was stored at −80°C in lysis RLT buffer (Qiagen) to which beta-mercaptoethanol had been added as per manufacturer’s guidelines for between 1 - 23 months.

### RNA extraction, cDNA processing and RNA sequencing

T cell samples underwent RNA extraction as per manufacturer’s protocol (Qiagen RNeasy kit) at CERA. All T-cell lysate samples, 135 GCA patient samples and 60 control samples, were randomised to RNA extraction batches of between 20–24 samples to avoid batch effects. RNA samples were eluted 30 μl in RNAse free water and stored at −80C until all extractions were complete. Samples were tested on the NanoDrop ND-100 spectrophotometer to check RNA quantity and quality (A260/A230 and A260/A280 between 1.8 and 2.1). Once all batches were extracted, samples were dispatched on dry ice to the Australian Translational Genomics Centre (ATGC) at Queensland University of Technology (QUT) for cDNA processing and RNA sequencing. At ATGC, RNA integrity (RIN) and quantity was confirmed with a Bioanalyzer 2100 (Agilent) before undergoing library preparation.

To avoid sequencing batch effects, all 195 samples (GCA *n=*135, and Control *n=*60) were re-randomised to be processed in one of three different cDNA library preparation batches (Illumina TruSeq Stranded mRNA Sample Preparation Kits). This kit purifies the polyA containing mRNA molecules. The Illumina Truseq protocol is optimized for 0.1−4 μg of total RNA and a RIN value ≥ 8 is recommended. The average total RNA yield varied between samples. The average RNA concentration was 137.9 ng/μl (range 12.1 to 1,130.0 ng/ul). Total RNA yield per sample averaged to 2,757.7 ng (range 242.0 to 22,600.0 ng) and average RIN was 8.9 (range 7.2 to 10.0). 600 ng total RNA was used to generate cDNA libraries (30 μl) for all samples with ≥600 ng total RNA available. Samples with less than 600 ng total RNA available were used entirely. Samples were barcoded to allow large throughput at sequencing. The number of PCR cycles for cDNA amplification was adjusted as required to equalise the cDNA yield as per the protocol. Quality control of library concentrations was assessed through LabChip GX High Sensitivity DNA assay.

RNA-Seq libraries were multiplexed and sequenced (75bp PE) in batches on an Illumina NextSeq500 high-throughput instrument. Each batch of cDNA libraries was pooled in equimolar volumes, and sequenced over three flow cells (FCs), with nine FCs used in total. To achieve uniform sequencing across a large number of samples, the data were reviewed following each run by determining the number of mapped reads per sample. The read count per sample volume pooled was used as a metric to re-pool the cDNA libraries for additional sequencing. As such the pool of cDNA libraries for each batch was adjusted so that all samples would reach 16M raw reads. This strategy also minimised between sample sequence run batch effects. cDNA libraries were sequenced and we obtained a median 11,017,433 mapped reads per sample and the read counts were aggregated into a single gene expression matrix. 40,744 transcripts had counts-per-million (cpm) > 1 in 50% of samples and underwent further analysis.

### Computational Analysis

Quality control of the sequencing data was performed on the FASTQ files. High quality reads were retained and Trimmomatic v0.36 was used to remove adapters and low quality bases. Reads were mapped to the GRCh38 human reference transcriptome using Kallisto v0.42.4 [10]. Only those with counts-per-million (cpm) > 1 in 50% of the samples were retained for further analysis. Transcript expression between libraries was normalised using the trimmed mean of M method (TMM) and corrected for batch effects using the *removeBatchEffect* function implemented in edgeR (Flowcell ID, Gender and Ethnicity) [11]. Hierarchical clustering and principal component analysis (PCA) confirmed the absence of batch effects and outlier samples (Supplementary Figure 2).

### Differential gene expression analysis

A total of 135 GCA samples (*n=*16 patients) spanning six timepoints and 60 control samples (*n=*16 patients) spanning two timepoints were grouped for analysis based on their CD4 (GCA=68, control=30) or CD8 MACS (GCA=67, control=30) separation. This grouping strategy formed the basis of the differential expression design matrix, allowing pairwise comparisons between individual timepoints on a case/control or CD4/CD8 basis. Differentially expressed transcripts were considered statistically significant if their false discovery rate (FDR) was less than 0.05. Differential expression (DGE) analysis between case and control subjects was performed comparing the initial T1 case specimens versus both the T1 and T2 of control specimens. Transcripts below FDR <0.05 and a two-fold change between cases and controls were considered significant.

### Polynomial modelling of transcript expression

The longitudinal expression profile of retained transcripts across six time points was tested for significant changes using polynomial regression. Polynomial regression modelling was performed with the patient weight-normalised steroid dosage fitted as a fixed effect. Steroid dose was normalised by dividing the Daily Steroid Dose by the Patient Weight. The global model *p-*value was corrected for multiple testing using the Benjamini-Hochberg method (FDR) and transcripts with an adjusted *p-*value below the FDR threshold (<0.05) were considered statistically significant.

### Functional enrichment and pathway analysis

Functional enrichment analysis was performed using the Reactome biological pathway database via the ReactomePA software package (version 1.18) and the CPdB web server (http://cpdb.molgen.mpg.de/) [12]. Pathway analysis results with adjusted *p-*values below the FDR threshold (< 0.1) were considered significant.

### Clinical phenotype regression analysis

Models were constructed to regress clinically relevant traits that were measured at the time of disease onset, or sample collection, against normalised gene expression levels. For quantitative clinical variables we used a linear model, and for categorical variables we used a logistic regression model. Clinical phenotypes were fitted against the expression of each of transcripts in GCA-only samples separated into CD4+ and CD8+ populations and weight-normalised daily steroid dose was included as a fixed effect. For each transcript, the adjusted *p-*value was calculated using the Benjamini-Hochberg method (FDR) method [13]. Transcripts with adjusted *p-*values below the FDR threshold (< 0.01) were retained for further analysis. The complete summary tables of tested phenotypes are available in Tables 2–4.

**Table 1:** Number of DE genes in each comparison

|  | CD4 |  | CD8 |  |
| --- | --- | --- | --- | --- |
| Contrast | DR | UR | DR | UR |
| Control 2 vs Control 1 | 0 | 0 | 0 | 0 |
| GCA T2 vs T1 | 0 | 0 | 0 | 0 |
| GCA T3 vs T1 | 1 | 8 | 35 | 80 |
| GCA T4 vs T1 | 2 | 7 | 1 | 3 |
| GCA T5 vs T1 | 0 | 0 | 0 | 0 |
| GCA T6 vs T1 | 0 | 0 | 2 | 0 |
| GCA T6 vs T3 | 0 | 0 | 45 | 10 |
| GCA T1 vs Control 1 | 67 | 129 | 93 | 188 |
| GCA T2 vs Control 1 | 254 | 228 | 325 | 453 |
| GCA T3 vs Control 1 | 196 | 190 | 1927 | 1783 |
| GCA T4 vs Control 1 | 179 | 200 | 576 | 827 |
| GCA T5 vs Control 1 | 1 | 1 | 101 | 296 |
| GCA T6 vs Control 1 | 0 | 0 | 1 | 1 |
| GCA T1 vs Control 2 | 22 | 58 | 58 | 156 |
| GCA T2 vs Control 2 | 276 | 233 | 187 | 335 |
| GCA T3 vs Control 2 | 194 | 171 | 1066 | 1227 |
| GCA T4 vs Control 2 | 197 | 179 | 351 | 615 |
| GCA T5 vs Control 2 | 2 | 0 | 55 | 222 |
| GCA T6 vs Control 2 | 0 | 0 | 0 | 0 |

**Table 2:** “Acute phase” symptoms, signs and relevant past medical history. Number of patients (total *n=*16) and genes significantly affected (FDR < 0.01) by clinical phenotype in regression models at T1.

|  | Phenotype | Number of patients with each feature at time of presentation | Number of Transcripts per cell type correlating to each phenotype |  |
| --- | --- | --- | --- | --- |
|  |  |  | CD4 | CD8 |
| 1 | Visual Disturbance | 14 | 23 | 247 |
| 2 | Temporal Headache | 14 | 67 | 34 |
| 3 | Other Headache | 13 | 30 | 76 |
| 4 | Scalp Tenderness | 12 | 10 | 7 |
| 5 | Malaise | 12 | 8 | 27 |
| 6 | Jaw Claudication | 11 | 70 | 10 |
| 7 | Fatigue | 11 | 6 | 29 |
| 8 | Loss of Appetite | 9 | 59 | 32 |
| 9 | Weight Loss | 8 | 27 | 55 |
| 10 | Fever | 4 | 177 | 41 |
| 11 | Polymyalgia Rheumatica | 4 | 51 | 53 |

**Table 3:**
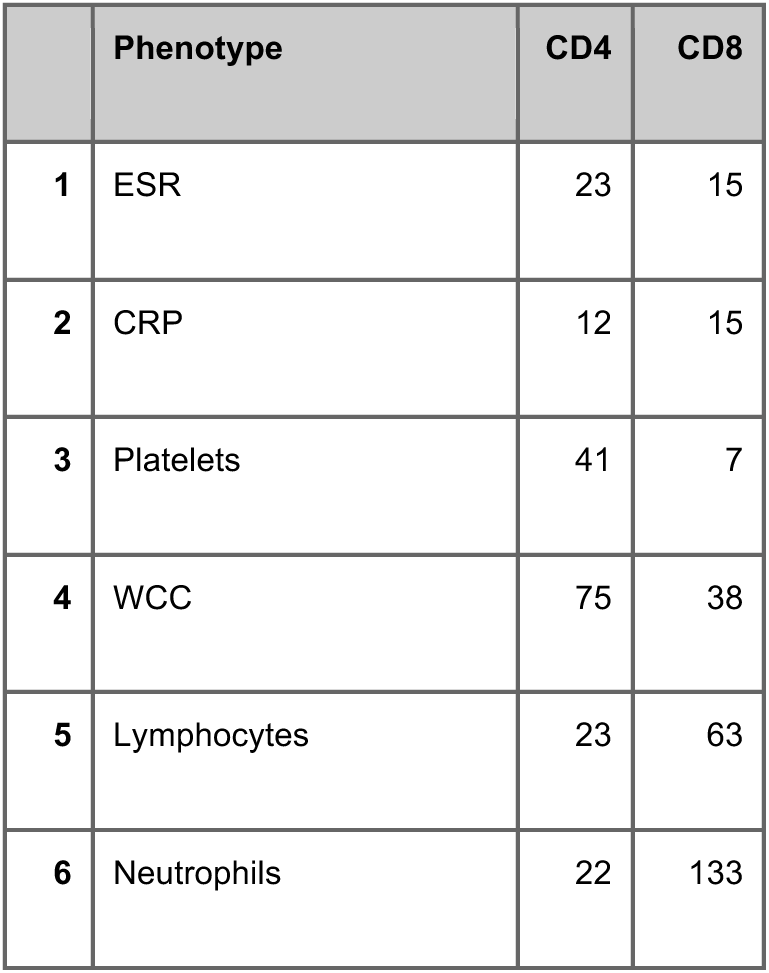
“Acute phase” biochemical markers. Number of genes significantly affected (FDR < 0.01) by biochemical markers in regression models at T1.

|  | Phenotype | CD4 | CD8 |
| --- | --- | --- | --- |
| 1 | ESR | 23 | 15 |
| 2 | CRP | 12 | 15 |
| 3 | Platelets | 41 | 7 |
| 4 | WCC | 75 | 38 |
| 5 | Lymphocytes | 23 | 63 |
| 6 | Neutrophils | 22 | 133 |

**Table 4:** “Prognostic genes”. Number of genes significantly affected (FDR < 0.01) by outcome and prognostic phenotype markers in regression models both in the acute phase alone (T1) as well as across all time points (T1-T6).

|  |  | T1 |  | T1-T6 |  |
| --- | --- | --- | --- | --- | --- |
|  | Phenotype | CD4 | CD8 | CD4 | CD8 |
| 1 | Monocular Blindness | 22 | 41 | 26 | 56 |
| 2 | Bilateral Blindness | 22 | 50 | 21 | 18 |
| 3 | Stroke/TIA | 40 | 4 | 153 | 70 |
| 4 | Relapse events | 6 | 3 | 47 | 166 |
| 5 | Deceased within 12 months | 878 | 904 | 43 | 50 |

## RESULTS

### Patient Recruitment and MACS events

16 incident patients with active GCA and 16 age-matched controls were recruited. The mean age was 78.2 years in the GCA cohort and 76.6 years in the control group. Both groups had the same 14:2 female to male ratio. Table 2 provides the number of patients presenting with the common symptoms and signs associated with GCA. Supplementary Tables 2 and 3 describe the specific ophthalmic manifestations and long-term prognoses observed in our patient cohort. Not all patients were able to complete 12 months of participation; therefore, not all patients had six samples collected (Supplementary Table 1). 6 patients were steroid-naive at T1; these patients had their first sample collected in the ED prior to commencing steroid treatment. Of the other 10 patients, 3 patients had been on steroids less than 24 hours, and the other 7 patients had been on steroids for between three to seven days at the time of T1.

In total, 195 MACS events (135 GCA and 60 control events) were performed, isolating between 2–10 million CD4+ and CD8+ cells per patient per time event. CD4+ MACS isolation resulted in greater cell counts than CD8+. The analysis on the CyAn ADP analyser shows good population purity after MACS-positive cell selection: an average of 97% for CD4+ cells and > 94% for CD8+ cells (Supplementary Figure 1).

### Differential expression analysis

To determine which transcripts showed the most variation in expression over the 12-month collection period, and to identify cell type specific signatures, we analysed the expression levels of samples from GCA patients (*n=*135) (Supplementary Figure 3). Figure 2 represents the expression levels of the top 40 most variable transcripts in CD4+ and CD8+ samples in GCA patients. The expression levels of control genes such as *CD4* and *CD8A/B* confirms the partitioning of CD4+ and CD8+ cells.

**Figure 2:**
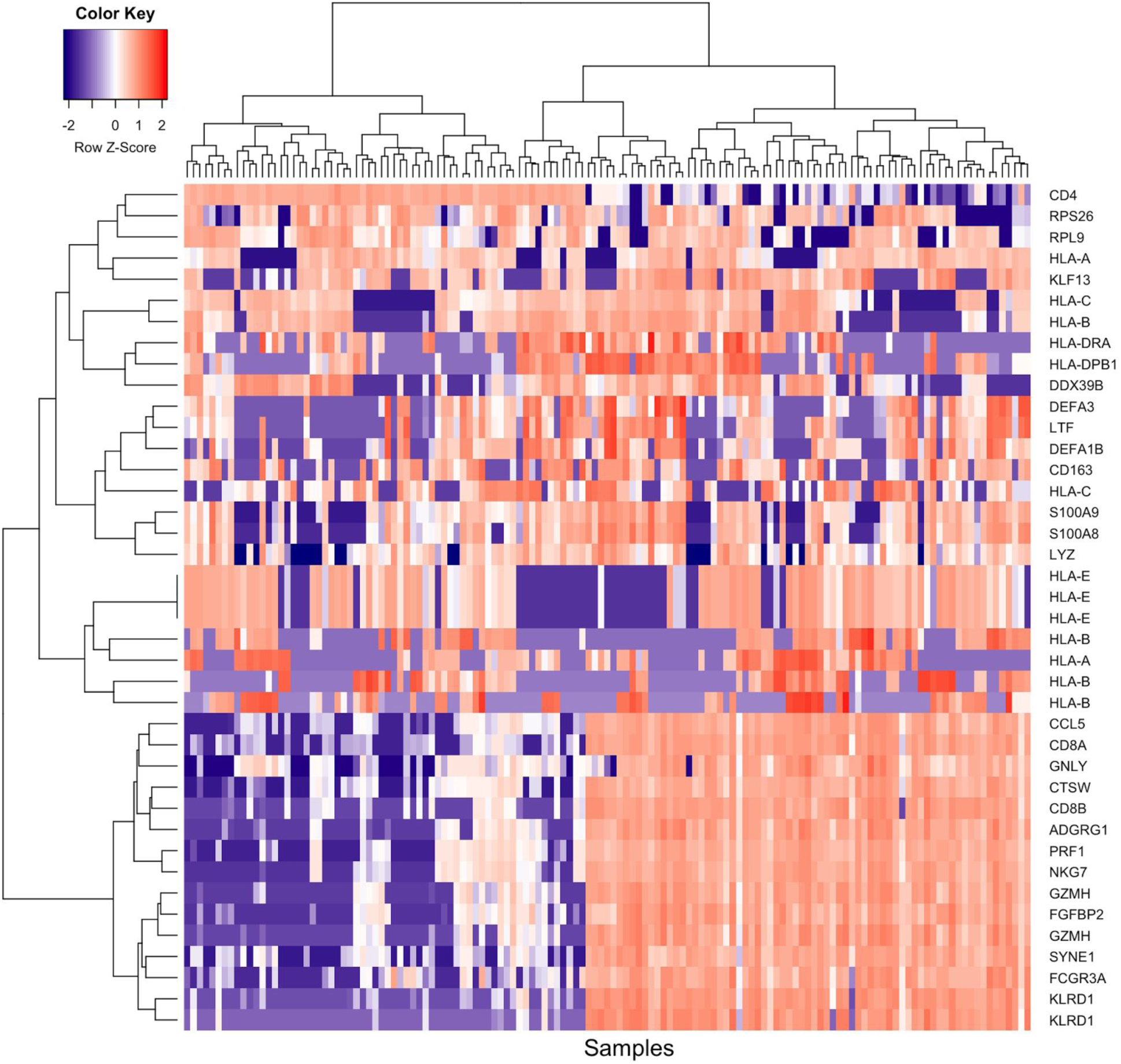
Expression levels of the top 40 genes with highest expression variation in CD4 and CD8 samples for all GCA patients. The color scale indicates normalised, log2-transformed gene expression (cpm), from low (blue) to high (red). Multiple gene IDs represent alternative transcript isoforms.

We investigated changes in gene expression in both CD4+ and CD8+ between cases and controls at T1. At a significance threshold of FDR < 0.05, we identified 67 down-regulated (DR) and 129 u*p-*regulated (UR) transcripts in CD4+ samples, and 93 DR and 188 UR transcripts in CD8+ samples (Table 1). The numbers of significantly differentially expressed transcripts increased dramatically at T3 in cases compared to the controls at T1 for CD8+ samples, and resolving to a near-control profile at T6. At T3 (6–8 weeks), we detected 1927 DR and 1,783 UR transcripts in CD8+ cells. Interestingly, DE transcripts in CD4+ cells reached a plateau from T2 to T4 (T2: 254 DR/228 UR; T3: 196 DR/190 UR; T4: 179 DR/200 UR).

We hypothesised that gene expression in GCA patients would return to baseline levels at approximately 12 months, corresponding to T6, marking disease quiescence. Transcripts remaining DE at T6 may be of clinical interest or mark evidence of previous disease despite current inactivity. In CD8+ cells, we identified two significant DE transcripts at T6 versus controls, *SGTB* (Small glutamine-rich tetratricopeptide repeat (TPR)-containing beta) and *FCGR3A* (Fc Fragment Of IgG Receptor IIIa), which showed log2 fold changes in expression of −0.54 (*p* = 4.83×10^−7^) and 1.99 (*p* = 1.75×10^−6^), respectively. There were no significant DE transcripts in the CD4+ cells between GCA T6 and the controls.

Differentially expressed genes between T1 and T6 in GCA patients could represent a biomarker of disease activity, marking either gene UR or DR during the acute phase of disease and then normalising as disease quiesces. From the CD8+ cell analysis, we detected two differentially expressed isoforms of *CD163* with significantly reduced expression levels. At T6 compared to T1, *CD163* isoform 1 (ENST00000359156) expression showed a log2 FC of −6.01 (*p* = 1.07×10^−6^), whereas the log2 FC of *CD163* isoform 2 (ENST00000432237) was −9.69 (*p* = 5.84×10^−8^). Notably, *CD163* expression is suppressed in response to pro-inflammatory stimuli in monocytes [14], and is inversely correlated with *CD16* expression [14,15], which is consistent with the increased *CD16* expression we observed in cases compared to controls at T6 (12 months). However, *CD16* was not consistently differentially expressed across all time points in CD8+ cells. There were no significant DE transcripts in the CD4+ cells between GCA T1 and T6. Reassuringly, no significant transcripts were observed in either CD4+ or CD8+ cells in the controls between T1 & T2. Tables of significant differentially expressed transcripts are presented in Supplementary Tables 4 (CD4) and 5 (CD8).

### Polynomial modelling of longitudinal transcript expression

To identify important transcripts whose expression levels vary across a 12-month period of the study, we used polynomial regression to model changes in the expression levels of 40,744 transcripts separately in CD4+ and CD8+ cells across the six timepoints. Using this approach, we detected 179 and 4 statistically significant expression profiles (FDR < 0.05) in CD4+ and CD8+ populations, respectively. Tables of significant transcript expression models are available in Supplementary Table 6.

The top 12 CD4+ profiles and all 4 significant CD8+ profiles are shown in Figure 3. In CD4+, the majority of genes demonstrated a pattern of decreased expression over the study course. Only two genes demonstrated a positive fold change and increase in expression levels over the 12 months, namely *FOXO1* involved in blood vessel development and *TRBC2* involved in complement cascade activation and phagocytosis. The four identified genes in CD8+ were *CCLN2, FANCA, PTCD2* and *THRAP3*. The first three genes demonstrate a negative log2 fold change, whilst *THRAP3* demonstrates an increased expression trend.

**Figure 3:**
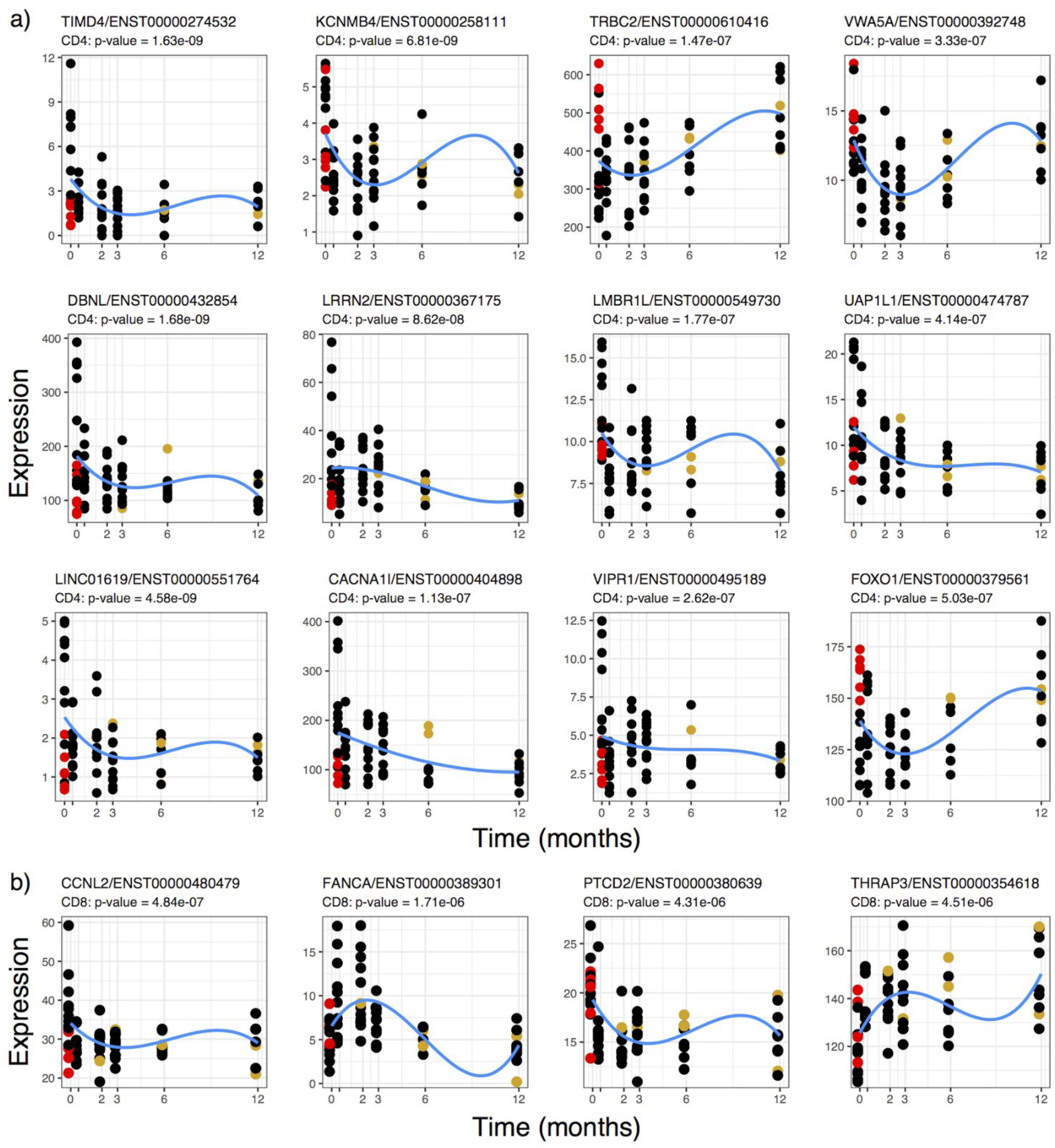
CD4+ cell **(a)** and CD8+ cell **(b)** polynomial regression. A polynomial model, with weight-normalised steroid dosage included as a fixed effect, was used to examine transcript expression over the duration of the study. Top transcripts with statistically significant expression profiles over the duration of the study are shown. The x-axis shows the duration of the study in months and the y-axis shows normalised expression levels (cpm). The red points represent the samples taken from steroid-naive individuals, and the gold points represent the samples taken from individuals who had suffered a relapse at the corresponding time point. The blue line shows the modelled expression values.

No substantial contribution of steroid dose to the model was observed across the 12-month time course (CD4: median beta = −0.001, median *p* = 0.439; CD8: median beta = −0.002, median *p* = 0.463). However, expression levels of certain genes at T1 may have been affected depending on whether patients were steroid-naive or had already been started on treatment at time of their first blood sample collection. Figures 3A and B highlight those patients who were steroid-naive in red and those who had already been started on steroid treatment in black. Expression of certain genes, for example *TIMD4, VIPR1*, and *FOXO1*, show obvious clustering depending on a patient’s treatment status and appear to be affected by corticosteroid initiation. Steroid treatment, even though only initiated in some instances less than 24 hours prior to blood collection at T1, has a clear effect on the expression of certain genes.

In CD4+, three genes, *LMBR1L, UAP1L1* and *KCNMB4*, showed least clustering at T1 and appeared least affected by steroid treatment, albeit having been through oral dose or intravenously administered prior to T1 collection. In CD8+ cells, *PTCD2* and *THRAP3* appear little affected by steroids at T1. *PTCD2* is highly expressed in both steroid-naive and patients on steroids at T1 and less so at T6, suggesting no major influence of steroids at T1. *THRAP3* shows increased expression over time suggesting that in the acute phase *THRAP3* expression might be suppressed.

From our DGE analysis, we observed significant reduction in *CD163* transcript expression between T1 and T6 in the CD8 cell population analysis. Our results for the polynomial expression modelling also reflected that *CD163* was significantly reduced at T6. However, model profiles of this transcript showed that the trend over the 12-month time course was not statistically significant (FDR > 0.05). Interestingly, we noted that several *CD163* isoforms in the analyses of both CD4+ and CD8+ cell populations had compelling model profiles. For all but one *CD163* isoform, expression levels returned to zero for all individuals at 12 months; however, these were not FDR-significant. The log2 fold-change in the expression of these transcripts over 12 months is shown in Figure 4.

**Figure 4.**
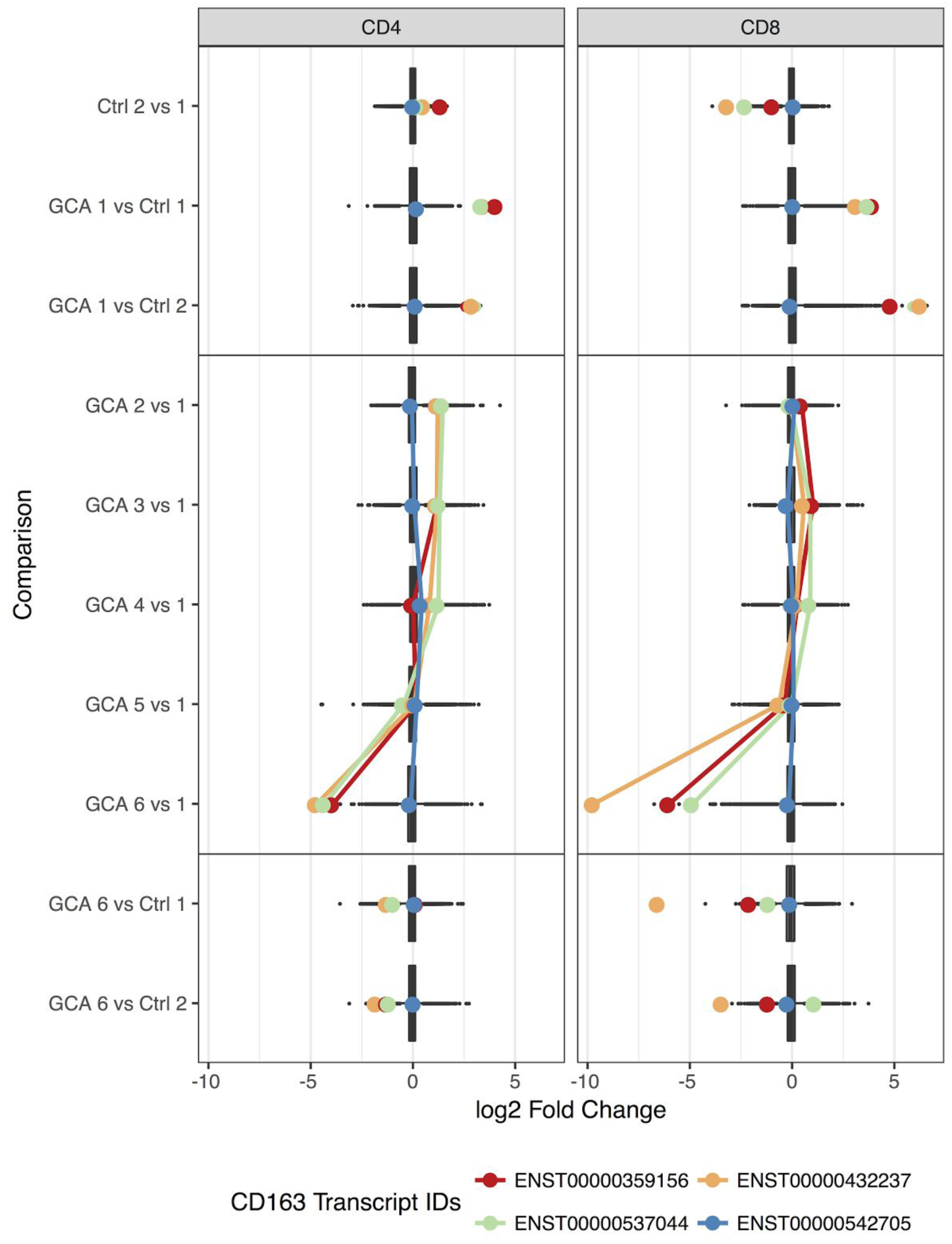
Fold-change distribution of differentially expressed transcripts in CD4 and CD8 samples for each differential expression comparison. Coloured points indicate the log2 fold-change of CD163 expression and shown for each transcript in CD4 and CD8 samples. Lines connect the fold-change values (log2-transformed) of differential expression comparisons along the time course only.

### Functional enrichment and pathway analysis

For individuals with GCA, we would expect an enrichment of immune and inflammation related pathways compared to healthy individuals. Biological pathway analysis of differentially expressed transcripts and statistically significant transcripts identified in the polynomial expression modelling analysis was performed using the curated Reactome database.

Significant DE transcripts in CD4+ samples comparing GCA to controls in the early time points showed a significant enrichment of T-cell receptor signaling (adj. *p-*value = 4.25 × 10^−3^; 11 genes). In CD8+ samples, we observed an enrichment of genes in pathways related to platelet degranulation (adj. *p-*value = 0.0124; 12 genes) and activation (adj. *p-*value = 0.0156; 20 genes), as well as Fc-gamma receptor (FCGR) dependent phagocytosis (adj. *p-*value = 0.0156; 13 genes). Furthermore, CD8+ samples from first two collected samples of GCA cases showed significant enrichment of pathways related to haemostasis (adj. *p-*value = 2.63 × 10^−6^; 118 genes), innate immune system (adj. *p-*value = 5.51 × 10^−6^; 169 genes) and the adaptive immune system (adj. *p-*value = 3.24 × 10^−4^; 129 genes).

Transcripts with a significant association across the 12-month collection time were interrogated for enrichment of specific biological pathways. We tested all 179 CD4 and 4 CD8 significant transcripts. In the CD4 transcripts, we observed an over-representation of transcripts in the integrin cell surface interactions (adj. *p-*value = 0.015) and Caspase-mediated cleavage of cytoskeletal proteins (adj. *p-*value = 0.0325) as well as cytokine signaling (adj. *p-*value = 0.08) and negative regulators of RIG-I/MDA5 signaling (adj. *p-*value = 0.08). In the CD8 results, there were insufficient significant transcripts to perform enrichment analyses. However, a literature search revealed *THRAP3* is involved in intracellular steroid hormone receptor signaling pathways, and *FANCA* in inflammatory responses and T-cell differentiation pathways.

### Clinical phenotype regression analysis

Linear and logistic regression models were used to estimate the effect of specific clinically important phenotypes on expressed transcripts. The analyses were three-fold. The first was to determine whether there were any genes that correlated with symptoms and signs used in the acute setting (T1) (Table 2). Second, we determined whether any genes directly correlated with the biochemical markers currently used in the acute phase (T1) (Table 3). Genes resulting from these first two analyses are potential biomarkers for disease activity in the acute setting and predict relapses. Thirdly, we determined gene correlations with markers of disease severity or prognosis (Table 4). These were categorised in terms of visual outcome: whether blinded in one eye, “monocular”, or both eyes, “bilateral”; relapse events; and whether the patient died during the study period. This enables us to identify genes that could provide prognostic information, ideally at the time of diagnosis (T1) but also during the course of disease (T1–6).

#### Correlation with clinical features in the acute setting

At the time of admission (T1), we would expect to observe some changes in gene expression to be strongly associated with clinical phenotypes related to the acute onset of disease. To identify a transcriptional signature that may be specific to active GCA, we examined the effect of clinically relevant phenotypes on gene expression in CD4 and CD8 samples taken at T1. Table 2 lists the eleven phenotypes and the number of statistically significant transcripts (FDR < 0.01) observed for each in CD4 or CD8 samples at T1. Genes or transcripts that are common to multiple symptoms/signs are likely to be clinically relevant, particularly at the acute onset of disease. In CD4 and CD8 samples, we identified 17 (CD4) and 27 (CD8) transcripts that were significantly associated with two or more clinical phenotypes.

In CD4 cells, *LAMTOR4* is a gene shared between jaw claudication and temporal headache, two important clinical features in acute GCA. Another gene associated with jaw claudication is *GZMB*, which is also associated with visual disturbance. *PPP1CB* and *EIF4A3* were shared by both jaw claudication and a background history of Polymyalgia Rheumatica (PMR). *EXTL3*, was expressed in both patients with jaw claudication and fatigue. We identified numerous genes associated with headache, both temporal and other types: *POFUT2* in CD4 cells, and *SLC35F6, HTD2, ZNF708, KLRC4-KLRK1* and *JMJD7* in CD8 cells. *EIF5A* in CD8 cells was common to both malaise and temporal headache. *SLA* and *ETS1* are genes shared by patients with a history of PMR diagnosis and those experiencing visual disturbances at T1.

Genes shared by three clinically important phenotypes at T1 are even more promising than those shared by two phenotypes and included 15 genes in CD4 and 16 in CD8 cells (Table 5). *SRRT* in CD4 was common to four phenotypes: death, fever, and both headache types. In CD8, *IL32* was common to five phenotypes: visual disturbance and raised neutrophils at T1, a history of PMR, and bilateral blindness and death within 12 months. The results for each phenotype are available in Supplementary Table 7.

**Table 5:** Genes associated with multiple phenotypes, both acute and prognostic, in CD4 and CD8 T cells.

| Gene | Phenotype 1 | Phenotype 2 | Phenotype 3 |
| --- | --- | --- | --- |
| **CD4** |  |  |  |
| *ATP1A1* | Temporal headache | Bilateral blindness | Death within 12 months |
| *LAMTOR4* | Temporal headache | Jaw claudication | Death within 12 months |
| *MATR3* | White cell count | Monocular blindness | Death within 12 months |
| *MLH1* | Temporal headache | Bilateral blindness | Death within 12 months |
| *NDEL1* | Loss of appetite | Other headache | Death within 12 months |
| *NDUFS7* | Temporal headache | Elevated lymphocytes | Death within 12 months |
| *PDZD4* | Fever | Loss of appetite | Reduced platelets |
| *POFUT2* | Temporal headache | Other headache | Death within 12 months |
| *RRP1* | Temporal headache | Bilateral blindness | Death within 12 months |
| *SDCCAG3* | Bilateral blindness | Relapse events | Death within 12 months |
| *SEC23A* | Fever | Reduced platelets | Death within 12 months |
| *SLC10A3* | Fever | Reduced white cell count | Death within 12 months |
| *USF2* | Temporal headache | Bilateral blindness | Death within 12 months |
| *WDR91* | Loss of appetite | Elevated white cell count | Death within 12 months |
| *ZNF343* | Scalp tenderness | Reduced neutrophils | Death within 12 months |
| **CD8** |  |  |  |
| *ACADVL* | Elevated neutrophils | Other headache | Death within 12 months |
| *CD6* | Elevated neutrophils | Visual disturbance | Death within 12 months |
| *EIF5A* | Malaise | Temporal Headache | Death within 12 months |
| *FDXR* | Loss of appetite | Weight loss | Death within 12 months |
| *INPPL1* | Malaise | Fatigue | Elevated neutrophils |
| *JMJD7* | Temporal headache | Other headache | Death within 12 months |
| *KIAA0513* | Visual disturbance | Bilateral blindness | Death within 12 months |
| *KLRC4-KLR1I* | Temporal headache | Other headache | Death within 12 months |
| *MTA1* | Elevated neutrophils | Visual disturbance | Death within 12 months |
| *NUCB2* | Temporal headache | Bilateral blindness | Death within 12 months |
| *PI4KA* | Elevated neutrophils | Visual disturbance | Death within 12 months |
| *PRAG1* | Elevated neutrophils | Bilateral blindness | Death within 12 months |
| *RNPS1* | Malaise | Fatigue | Death within 12 months |
| *SLC35F6* | Temporal headache | Other headache | Death within 12 months |
| *UQCRC1* | Malaise | Other headache | Death within 12 months |
| *ZNF708* | Temporal headache | Other headache | Death within 12 months |

#### Correlation with currently used biochemical markers

We asked whether the results of several routine blood tests, including white cell count, platelet count, ESR and CRP correlated with changes in gene expression (Table 3). We observed significant clinical associations for each biochemical marker in both CD4 and CD8 samples.

Thrombocytosis - raised platelet count - is a good predictor of acute GCA [16]. Our analysis revealed associations of multiple genes common to both raised platelet count and fever in CD4 cells, namely *ATP9B, SEC23A, PDZD4, ABCA2, ELK1, CCDC88C* and *DGKZ*. In addition, ESR and CRP are biomarkers commonly used to predict the likelihood of GCA, and we found that *SAP18* in CD4 was associated with raised ESR and jaw claudication, whereas in CD8 cells *AMPD2* was associated with raised CRP and visual disturbances.

White-blood cell count (WCC), neutrophil and lymphocyte count may also be affected in GCA, although this may be due to the corticosteroid treatment rather than the inflammatory process [17]. In the CD4 cells of our patients, we found that *SPPL2B* expression was common to both those with raised WCC and jaw claudication whilst *MATR3* was associated with raised WCC and long-term monocular blindness. *NDUFS7* expression in CD4 cells was associated with an increased lymphocyte count and temporal headache in CD4, whereas in CD8 cells *AP1G2* was common to raised lymphocytes and visual disturbance. Additionally, expression of *ZNF343* and *INTS14* in CD4 cells were associated with both raised neutrophil and with scalp tenderness and event relapses respectively.

#### Correlation with prognostic outcome 12 months after diagnosis

We identified genes that overlap between phenotypes marking acute disease as well as those marking prognosis. For example temporal headache at T1 as well as bilateral blindness showed significant association with CD8 expression of *TCF7* (*TH:* beta = −0.151, adj. *p-*value = 6.0 × 10^−4^, BB: beta = −1.801, adj. *p-*value = 2.2 × 10^−3^) and *NUCB2* (beta = 1.571, adj. *p-*value = 1.31 × 10^−6^). The expression of such genes could provide insight into visual prognosis in those patients presenting with headache in GCA. *RPL17* in CD8 was associated between jaw claudication and relapse events, and *FTSJ1* in CD4 between jaw claudication and long-term cerebrovascular events. Many genes were shared between multiple acute phase phenotypes and mortality within 12 months (Table 5). Figure 5 shows the network analysis of clinically correlated phenotypes with shared genes, and highlights the link between phenotypes through significant shared genes.

**Figure 5.**
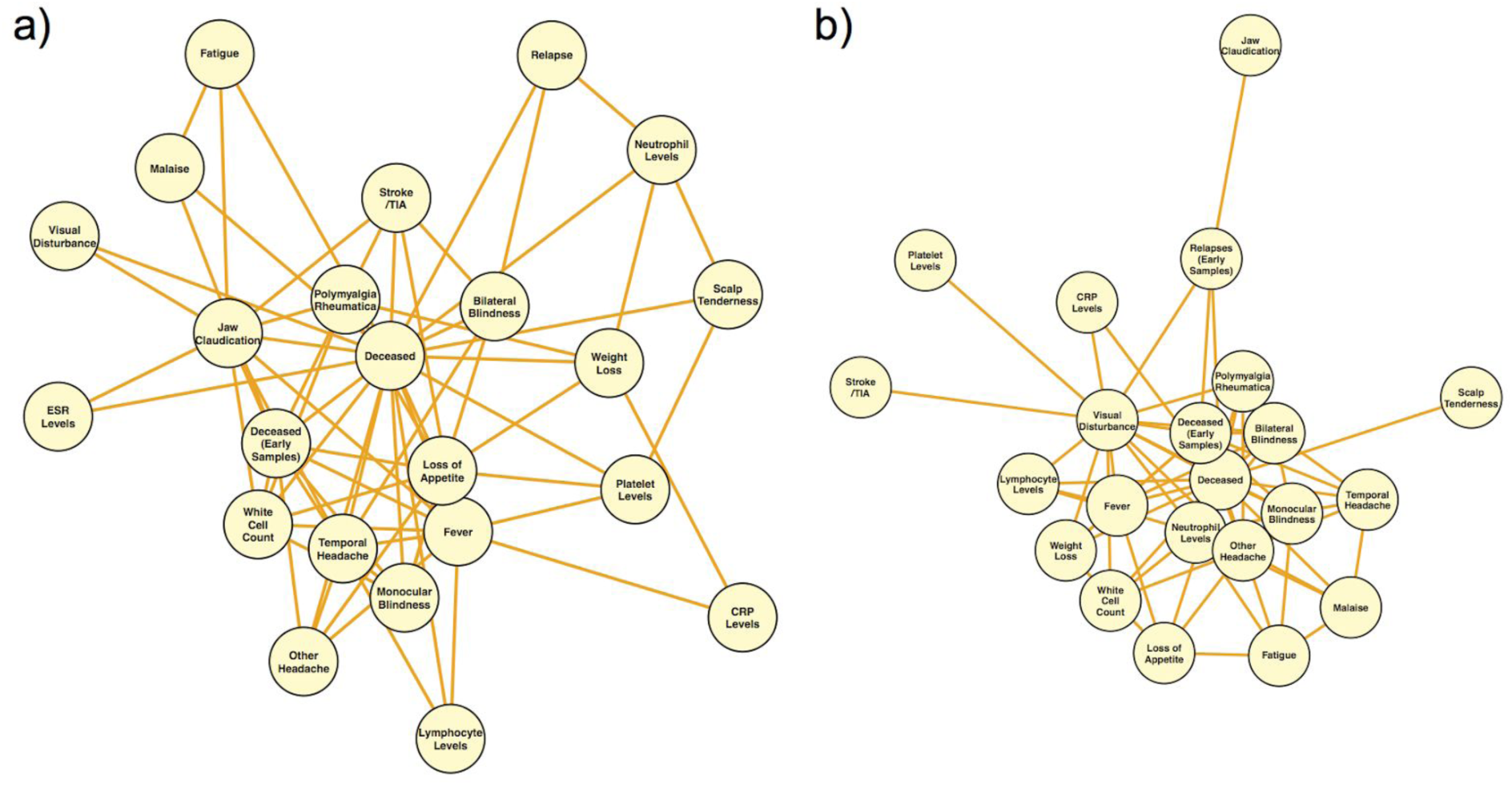
Network analysis of clinically correlated phenotypes with shared genes. Network plots show the clinical phenotypes observed for GCA patients at the time of presentation with shared, statistically significant genes (FDR < 0.01) in **(a)** CD4 and **(b)** CD8 samples. Each network node represents a phenotype that shares significant genes with > 1 other phenotype. Network edges represent connections (shared genes) between phenotypes.

## DISCUSSION

Through transcriptional profiling of T-lymphocyte we identified 4,031 genes in CD4+ and CD8+ cells (CD4: 884; CD8: 3,147) that are differentially expressed between patients with active GCA compared to age- and sex-matched controls. Longitudinal profiling of cases was undertaken with the aim of distinguishing genes that are up- or down-regulated during the acute phase of disease, which later normalise as the disease quiesces. We hypothesised that gene expression in GCA patients would return to normal at approximately 12 months. With polynomial modeling analysis of the significant differentially expressed genes, we identified 4 transcripts in CD8+ cells and 179 in CD4+ cells that show a change in expression profile over the course of twelve months (Figure 2). As there were no statistically significant differentially expressed genes between both samples taken from controls subjects at separate times, the genes we report as differentially expressed likely represent true changes occurring in GCA disease activity.

Next, we determined whether the fold change in expression was secondary to the true effect of disease status rather than due to steroid treatment. It is important to take into consideration steroid influence on gene expression, especially early in the treatment course, as this would allow for the identification of a biomarker that could help diagnose GCA in the acute setting prior to treatment. As patients received high-dose corticosteroids between T2-T6, we compared gene expression of those patients who were steroid naive versus those who had already been initiated on treatment at their first sample collection. *LMBR1L, UAP1L1* and *KCNMB4* in CD4, and *PTCD2 and THRAP3* in CD8, showed least clustering at the initial collection and seemed least affected by steroids at T1 (Figure 3), suggesting that the expression profiles of these genes seen in patients, compared to controls, is likely representative of “acute disease” at T1 rather than a steroid-induced change.

Gene expression patterns seen from our polynomial modeling analysis over the 12 months might have been influenced by systemic corticosteroid treatment (Figure 3). In CD8+ samples, differential expression of certain genes increased dramatically at around 6–8 weeks (T3) in cases compared to the controls, and in CD4+ cells, differential expression plateaued from T2-T4. Duration of steroid treatment did not have a significant effect on expression and was removed from analysis. We also adjusted for steroid dose and patient weight in our analysis; however, the peak in expression in both cell types at these time points could be caused by a delayed or accumulation of steroid-induced effect. Nevertheless, from a diagnostic perspective, acute phase evaluation at T1 is most crucial for patient assessment and this potential delayed steroid-induced effect is not that problematic in our analysis. It does, however, make evaluation of expression levels in relation to relapse events between 0.5–12 months (T2-T6) slightly challenging.

Our results show that transcripts that remain DE at 12 months (T6) could potentially be used in clinical practice to detect evidence of previous GCA disease despite current inactivity. In CD8+ cells, we identified two significant differentially expressed transcripts at T6 versus controls, *SGTB* and *FCGR3A*. Little is known about *SGTB* but it has been associated with neuronal apoptosis after neuroinflammation [18]. Interestingly, *FCGR3A* encodes CD16a, which forms part of the Fc receptor of the immunoglobulin complex and interacts with a number of immune-related proteins including CD4 and PTPRC, a protein required for T-cell activation. Recently, Lassaunière et al. showed that Black individuals have significantly reduced proportions of *FCGR3A* natural killer cells (95.2% vs. 96.9%) and CD8+ T lymphocytes (9.6% vs. 11.7%) compared to Caucasians [19], and this may serve as a predictive marker for a high-expressing *FCGR3A* phenotype in Caucasians, the population most affected by GCA. A recent genome-wide association study revealed that the FCGR2A/FCGR3A genes confer susceptibility to Takayasu arteritis, another chronic large-vessel vasculitis [20]. Furthermore, two recent studies investigating rejection in heart and kidney transplants, observed selective changes in endothelial/angiogenesis and natural killer cell transcripts, including *CD16A* and *FCGR3A* which showed increased expression with rejection phenotypes [21][22]. Both studies illustrate the clinical potential of gene transcripts to illustrate transplant rejection diagnosis. A future study would need to be conducted to investigate the expression of *FCGR3A* and *CD16a* at the arterial level (TAB) of GCA patients to determine whether increased expression at local level is representative to that found in peripheral T-cells. If so, *FCGR3A* could potentially be used as a biomarker of GCA severity in peripheral blood.

From our CD8+ cell analysis, we detected two differentially expressed isoforms of *CD163* with significantly reduced expression levels at first and last collection points. *CD163*, however, is a member of the scavenger receptor cysteine-rich (SRCR) superfamily, and is mostly expressed in monocytes and macrophages [23]. Despite an excellent T-cell population purity of >97% isolated through MACS (Supp Fig 1), monocytes and macrophages may carry CD4+ and CD8+ cell surface markers as T lymphocytes, and may have carried over into our final positively-selected T-cell population. Irrespective of its derivative cell population, *CD163* expression may play a crucial role in the context of GCA and, as a result, provide crucial information. *CD163* is involved in dendritic cell development, a cell crucial in the pathogenesis of GCA [24]. It has been suggested that the soluble form of *CD163* (sCD163) may have an anti-inflammatory role, and be a valuable diagnostic parameter for monitoring macrophage activation in inflammatory conditions where macrophage function is affected [25]. A number of clinical studies have evaluated the role sCD163 as a disease marker in inflammatory conditions including autoimmune disease, transplantation and cancer [26][27][28]. Expression levels of *CD163* were reduced in our patients at T6, possibly reflecting disease quiescence. It is likely that 12 months after disease onset, the need for CD163-monocytes and macrophages to clear damaged tissue has become redundant. *CD163* featured in both our differential expression and polynomial regression analyses and therefore warrants further investigation in the context of GCA, potentially through study of peripheral or tissue monocytes and macrophages.

Another strength of this study is that, through linear and logistic regression analyses, we identified associations between specific clinically important phenotypes and expressed transcripts. We detected genes which correlated with both symptoms and signs as well as biochemical markers used in the acute setting (Table 2). Symptoms causing the most suspicion of a potential GCA diagnosis consist of jaw claudication, temporal headache (or other type), scalp tenderness and visual disturbance [1]. Genes shared by multiple of these phenotypes are likely to be particularly relevant to making a diagnosis and could be used as biomarkers for disease activity in the acute setting and potentially predict relapses.

Jaw claudication is often considered the most predictive symptom of GCA; for example, a patient has a nine time greater risk of a positive TAB when they experience jaw claudication [29]. In CD4 cells of our patient cohort, *LAMTOR4*, was shared between jaw claudication and temporal headache. This protein is part of the ragulator complex, which is involved in pathways regulating cell size and cell cycle arrest [30]. A gene common to both jaw claudication and visual disturbance is *GZMB*, otherwise known as Granzyme B enzyme. *GZMB* is necessary for targeting cell lysis in cell-mediated immune responses and is involved in the activation of cytokine release and cascade of caspases responsible for apoptosis execution. Its involvement has been reported in other autoimmune diseases such as type 1 diabetes and systemic lupus erythematosus [31,32]. *PPP1CB*, linked to vascular smooth muscle contraction pathway [33], was common to patients with jaw claudication and a background history of PMR, which has been shown to increase the risk of GCA [34]. *EXTL3*, involved in the heparan sulfate biosynthesis pathway and previously associated with syphilis, was expressed in both patients with jaw claudication and fatigue [35].

Multiple genes were associated with temporal and other types of headache in our patients. These included *POFUT2* in CD4 cells and *SLC35F6, HTD2, ZNF708, KLRC4-KLRK1* and *JMJD7* in CD8 cells. These genes have been described as involved in cellular defense mechanism, innate immunity, cell proliferation and apoptosis signaling pathways [36]. One example of great clinical interest is a gene shared by patients with a history of PMR and those experiencing visual disturbances at T1. *ETS1*, controls lymphocyte differentiation, modulates cytokine and chemokine expression. Low expression levels of *ETS1*, leading to aberrant lymphocyte differentiation, have been found in systemic lupus erythematosus [37]. *ETS1* also has a potential role in the regulation of angiogenesis [38]. *ETS1* warrants further functional investigation in relation to its vascular role and as a biomarker for GCA for those patients presenting with PMR.

We determined gene correlations with markers of disease prognosis and severity (Table 3). *G*enes in association with poor prognostic outcome markers of GCA, such as blindness, relapses and death could provide useful predictions in the acute setting and could help determine the treatment intensity and length required for those particular patients. We identified genes that overlap between acute phase markers as well as the prognostic markers. For examples temporal headache at T1 as well as bilateral blindness showed significant association with CD8 expression of *TCF7*, which is important for adaptive T lymphocyte and innate lymphoid cell regulation [39]. Both these phenotypes were also associated with *NUCB2*, which encodes Nesfatin-1. *NUCB2* is linked to inflammation and coagulopathies, and is correlated with mortality following brain injury [40]. As *TCF7* and *NUCB2* expression are associated with temporal headache in patients with GCA, these genes could also raise suspicion of poor visual outcome in patients presenting with temporal headache with GCA diagnosis.

We identified 15 genes shared across three phenotypes in CD4 and 16 across CD8 cells (Table 5). In CD4 cells, *SRRT, a gene* associated with cell proliferation [41], was common to four phenotypes: death, fever, and both types of headaches. In CD8, *IL32*, a member of the cytokine family [42], was common to 5 phenotypes: a history of PMR, visual disturbance and raised neutrophils at T1, bilateral blindness and death within 12 months. *IL-32* involvement has been described in vasculitides such as granulomatosis with polyangiitis and anti-neutrophil cytoplasm antibodies (ANCA) associated vasculitis ([43,44]. A previous quantitative gene expression analysis study investigating *IL-32* in GCA demonstrated a strong and significant u*p-*regulation of *IL-32* in TAB specimens of patients with GCA; in particular it was highly expressed by vascular smooth muscle cells of inflamed arteries and neovessels within inflammatory infiltrates [45]. This study also evaluated circulating CD4+ Th1 lymphocytes by flow cytometry which showed that there was a greater abundance of them in GCA patients than controls and that they produced greater amounts of *IL-32* [45]. From our study, expression of *IL32* in patients presenting with visual disturbance, a history of PMR in the presence of an abnormal neutrophil count, should raise suspicion of GCA diagnosis with poor prognostic outcome. Altered expression of these genes should raise suspicion of GCA diagnosis with poor outcome. Such genes warrant more investigation in the context of GCA as these correlated with not only clinical and biochemical phenotypes but also with prognoses.

GCA is a devastating disease associated with significant morbidity and mortality. The current mainstay treatment of high-dose corticosteroids is effective but is commonly associated with potentially serious complications affecting up to 89% of those with GCA[3]. Even after successful initial treatment with corticosteroids, GCA relapses in up to two-thirds of patients[46]. As shown by our study, 5 out of 16 patients experienced relapses requiring an increase in steroid dose (Supplementary Table 1). Unlike in other autoimmune diseases, most steroid-sparing agents and the use of adjunct agents in GCA_[MB1]_ are not associated with a significant improvement in outcome[46,47]. Tocilizumab, a humanized monoclonal antibody directed against the IL-6 receptor, has been found to improve both induction and maintenance of remission in patients with GCA for up to 12 months [48]. However, there is a large side effect profile from toxilizumab. Interestingly we did not see DGE for IL-6.

In summary, this study has identified genes potentially implicated in the patho-aetiology of GCA that could be used as biomarkers to monitor disease activity and to predict outcome. Further functional investigation is needed to understand the pathways in which these genes play a role in the pathogenesis of GCA and also to determine whether the DGE in this study can be translated into the clinical setting as new potential biomarkers and assist in finding more effective and safer treatments for GCA.

---

Supplementary Figures
1. FACS
2. Batch correction (SuppFig2.png)
3. Top 500 variable genes by group (CD4/CD8) (SuppFig3.png)

Supplementary Tables
1. Cases recruited
2. Ophthalmic clinical summary data
3. General Disease Outcome and prognostic measures
4. DE results CD4 (file: SuppTable4_DE_CD4.xlsx)
5. DE results CD8 (file: SuppTable5_DE_CD8.xlsx)
6. Significant polynomial modelling results (file: SuppTable6_PolynomialModelling_CD4CD8.xlsx)
7. Significant clinical correlation data package (file: SuppTable7_sig_clinical_correlations.zip)

a. Shared gene overlap
b. DE gene overlap
c. Significant genes per phenotype
d. Tables of results for each individual correlation

**Supplementary Figure 1.**
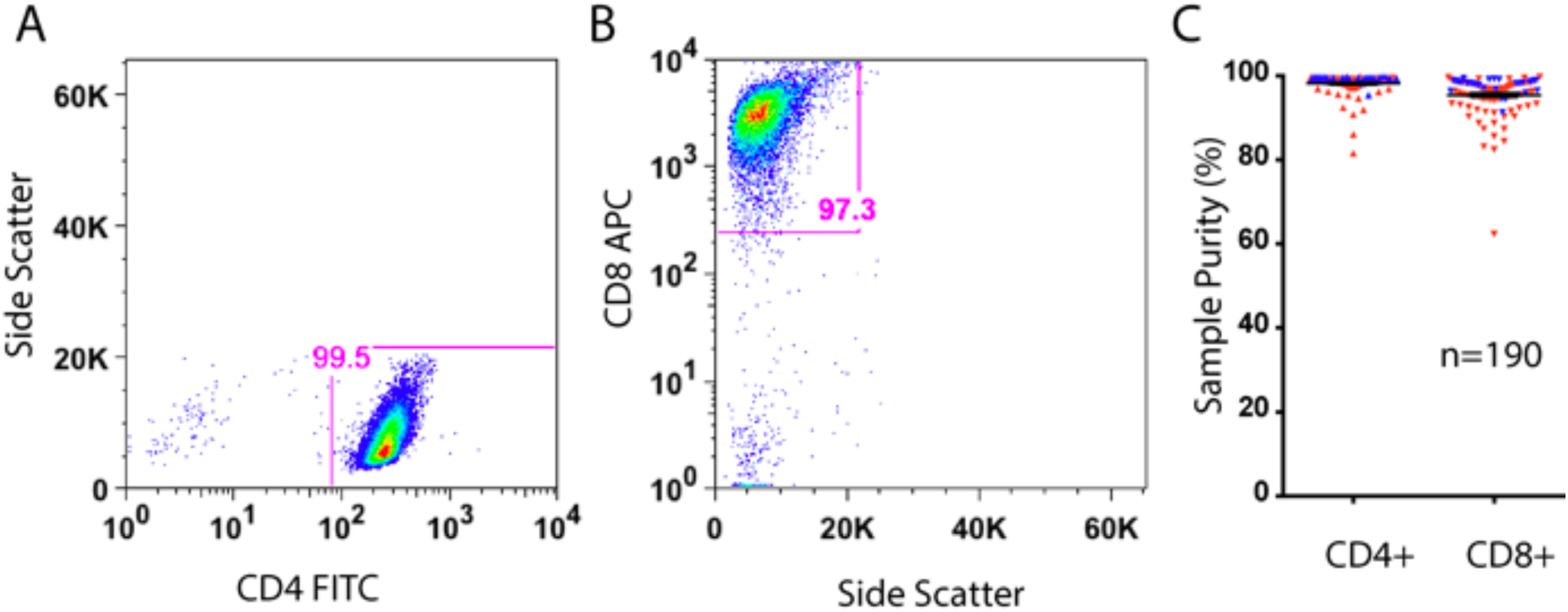
Quality control metrics for stored specimens. Representative FACS analysis for FITC bound CD4 (**A**) and APC bound CD8 cells (**B**). Panel **C** displays the FACS confirmed purity of all specimens, with case and control samples represented by red and blue triangles respectively.

**Supplementary Figure 2.**
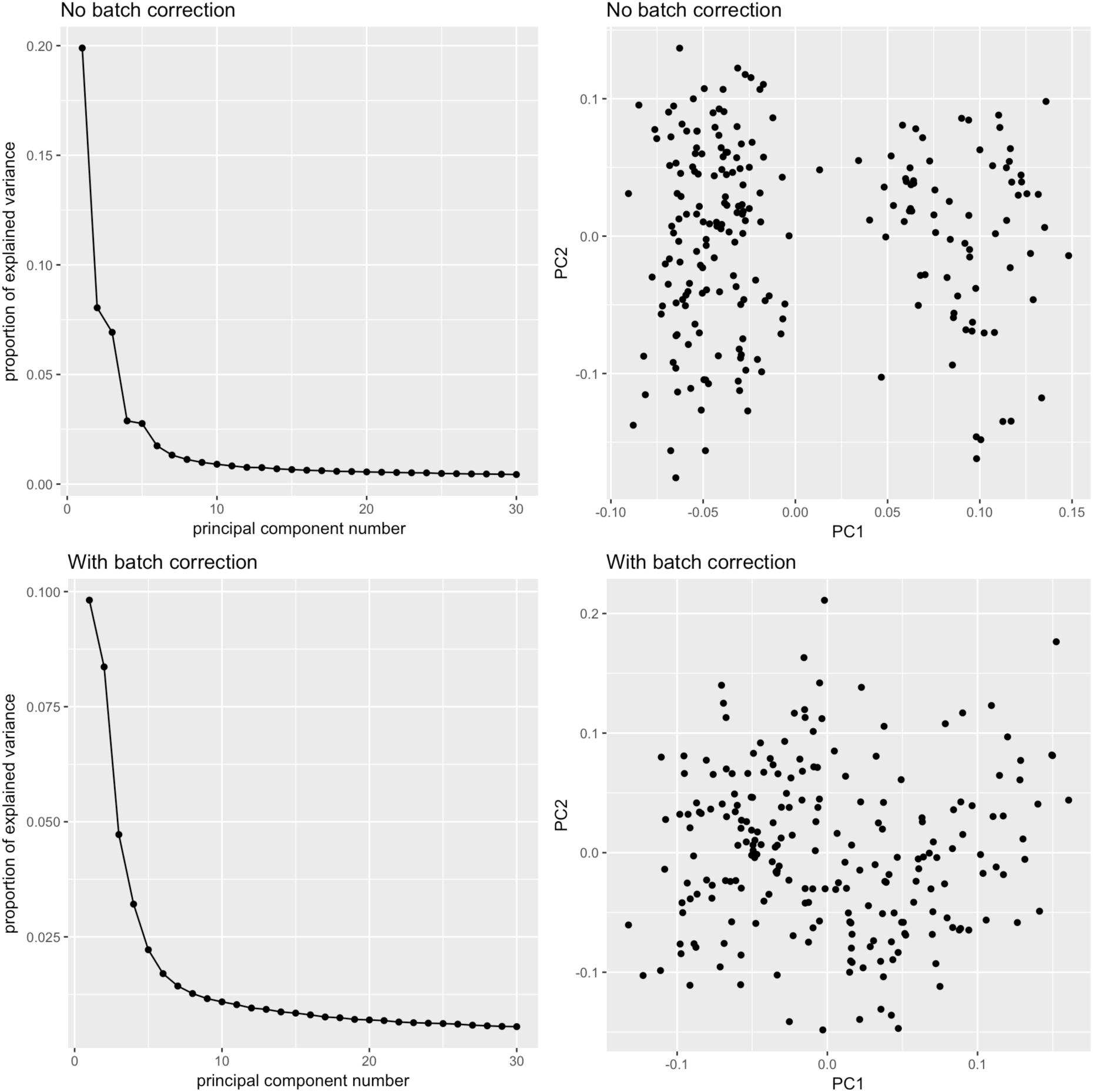
Effect of batch correction on 195 samples (2 samples of the 197 were removed). Three parameters (Flowcell ID, Gender and Ethnicity) were used to remove confounding effects in edgeR. PC1 contributes the greatest amount of variance and is largely attributed to Flowcell ID, which accounts for most of the variance in sequencing experiments.

**Supplementary Figure 3.**
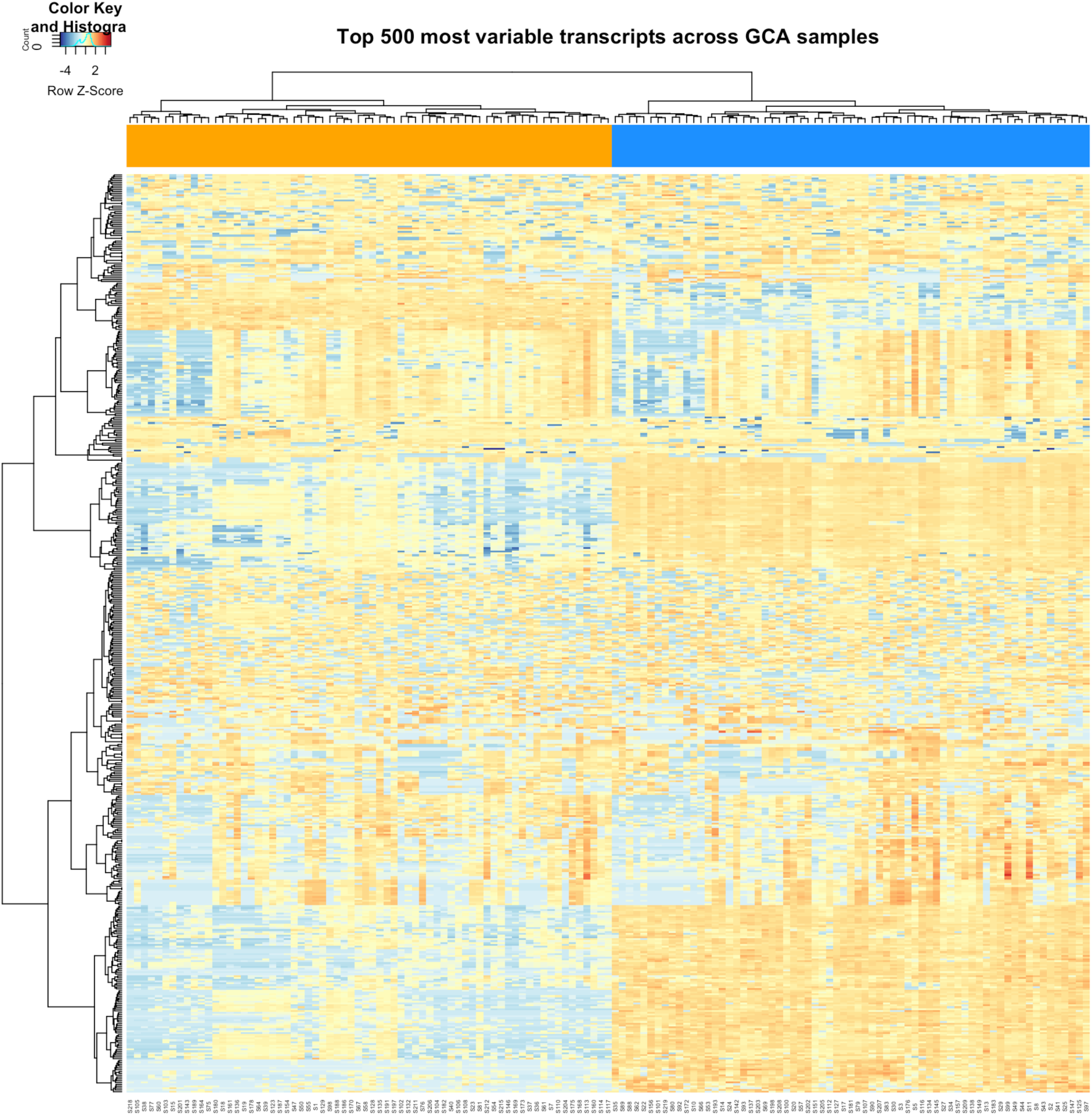
Expression levels of the top 500 most variable transcripts in CD4 and CD8 cells, shown for each of 135 samples. Sample groups are indicated by the orange (CD4) and blue (CD8) bars at the top of the heatmap.

**Supplementary Table 1.** Cases recruited and attendance for all 6 time points. Abbreviations: T1 (Day 0–7); T2 (2–3 weeks); T4 (~3 months); T5 (~6 months); T6 (~12 months); DNA, did not attend.

| GCA case | T1 | T2 | T3 | T4 | T5 | T6 |
| --- | --- | --- | --- | --- | --- | --- |
| 1. | 0 | 17 | 42 | 103 | 189 | 374 |
| 2. | 5 | 17 | 54 | 99 | 187 | 383 |
| 3. | 6 | 18 | 34 | 89 | 194 | 377 |
| 4. | 1 | 29 | DNA | 99 | 190 | 386 |
| 5. | 1 | 24 | 51 | 80 | 164 | 390 |
| 6. | 6 | 18 | 34 | 83 | 175 | 369 |
| 7. | 3 | 21 | 51 | 93 | 189 | 387 |
| 8. | 0 | 7 | 49 | 90 | End of study |  |
| 9. | 0 | 11 | 49 | Deceased |  |  |
| 10. | 0 | 7 | 35 | 90 | End of study |  |
| 11. | 0 | DNA | 35 | 91 | 182 | 386 |
| 12. | 2 | Absent due to illness |  |  |  | 366 |
| 13. | 3 | 19 | 55 | 90 | Withdrew from study |  |
| 14. | 1 | 17 | 65 | Deceased |  |  |
| 15. | 5 | Deceased |  |  |  |  |
| 16. | 0 | DNA |  | 89 | End of study |  |
| Mean | 2.0 | 17.1 | 46.2 | 91.5 | 183.75 | 379.8 |
| **Median** | 1 | 17.5 | 49 | 90 | 188 | 383 |
| **(Range)** | (0-6) | (7-29) | (34-65) | (80-103) | (164-194) | (366-390) |

**Supplementary Table 2.** Ophthalmic clinical summary data (GCA cases, n=16).

|  | Number of cases |
| --- | --- |
| ***Visual disturbance at T1*** |  |
| Monocular | 10 |
| Bilateral | 4 |
| None | 2 |
| ***Eye affected*** |  |
| Right | 4 |
| Left | 5 |
| Both | 4 |
| ***Ophthalmic Manifestation*** |  |
| Amaurosis Fugax | 2 |
| AION | 9 |
| Transient diplopia | 1 |
| 3rd Cranial nerve palsy | 2 |
| ***Final recorded visual outcome*** |  |
| Normal vision both eyes | 4 |
| Normal vision one eye, visual impairment other | 7 |
| Blind one eye | 2 |
| Blind both eyes | 3 |

Definitions (Snellen Chart): *Normal vision*: >6/9; *Vision impairment*: <6/9 but >6/60; *Severe vision impairment ("Blind")*: <=6/60.

**Supplementary Table 3.** General Disease Outcome and prognostic measures

| Category | Number of cases |
| --- | --- |
| *Number of relapse events during the 12 month study period* |  |
| None | 5 |
| One | 5 |
| Two | 2 |
| Three or more | 1 |
| Unknown (loss to follow up) | 3 |
| *Deceased within 12 months* |  |
| Yes | 3 |

